# Molecular Phenotyping of Oxidative Stress in Diabetes Mellitus with Point-of-care NMR system

**DOI:** 10.1101/565325

**Authors:** Weng Kung Peng, Lan Chen, Bernhard O Boehm, Jongyoon Han, Tze Ping Loh

## Abstract

Diabetes mellitus is one of the fastest growing health burdens globally. Oxidative stress which has been implicated to the pathogenesis of diabetes complication (e.g., cardiovascular event) were, however, poorly understood. We report a novel approach to rapidly manipulate the redox chemistry (in a single drop) of blood using point-of-care NMR system. We exploit the fact that oxidative stress changes the subtle molecular motion of water-proton in the blood, and thus inducing a measurable shift in magnetic resonance relaxation properties. This technique is label-free and the whole assays finish in a few minutes. Various redox states of the hemoglobin were mapped out using our newly proposed two-dimensional map, known as T_1_-T_2_ magnetic state diagram. We demonstrated the clinical utilities of this technique to rapidly sub-stratify diabetes subjects based on their oxidative status (in conjunction to the traditional glycemic level), to improve the patient risk stratification and thus the overall outcome of clinical diabetes care and management. (155 words)

**Key Points for Summaries:**

1. A novel approach to rapidly manipulate the redox chemistry (in a single drop) of blood using point-of-care NMR system.
2. Assessment of the oxidative status, in conjunction to their glycemic level allows sub-stratification of diabetes subjects which was demonstrated clinically.

**Visual Abstract:** 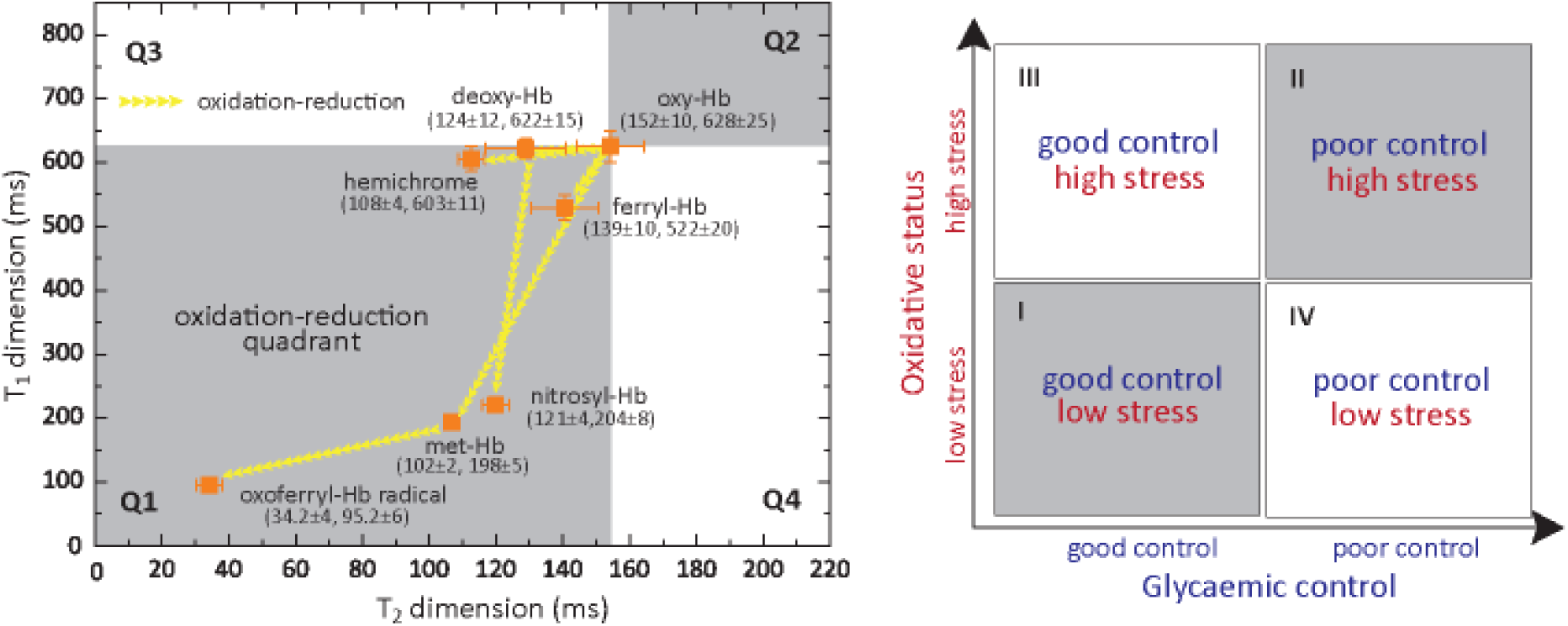

## INTRODUCTION

Diabetes mellitus (DM) is one of the fastest growing health burdens that is projected to affect 592 million people worldwide by 2035^1^. DM is defined by a persistent elevation of plasma glucose concentration. Under chronic hyperglycemic condition, glucose is non-enzymatically attached to protein (glycation), which has deleterious effects on their structure and function. Hence, glycated hemoglobin A_1c_ (HbA_1c_), which reflects the overall glycemic burden of an individual over the previous 2─3 months, is increasingly used to diagnose the disease^2^. It is also recommended for monitoring long-term glucose control of DM patients, and for risk stratification^3, 4^.

However, HbA_1c_ does not adequately reflect all the disease associated risk factors. In particular, restoring HbA_1c_ level to near-normal level does not necessarily translate into a significant reduction of cardiovascular event, a diabetes complication commonly associated with oxidative stress^5^. In addition, subjects with stable chronic hyperglycemia due to glucokinase mutations were found to have unexpectedly lower prevalence of micro/macrovascular complication. A major pathological effect of diabetes mellitus is the chronic oxidative—nitrosative stress and recently reported carbonyl^6^ and methylglyoxal stress^7^, which drives many of the secondary complications of diabetes including nephropathy, retinopathy, neuropathy, and cardiovascular diseases^8^. Oxidative-nitrosative stress can damage nucleic acids, lipids and proteins, which severely compromise the cellular health and induce a range of cellular responses leading ultimately to cell death^9–11^. Direct measurement of oxidative stress and susceptibility in patients may improve the prediction of disease associated risks related to oxidative stress, and hence improve the long term diabetes care and management program^12, 13^.

Currently, an individual’s oxidative status cannot be easily characterized in detail using routinely available biomarkers^14^ in clinical practice and/or at point of care. This has impeded the understanding of the pathological effects of acute and prolonged exposure to oxidative stress. The reactive oxygen species (ROS) and reactive nitrogen species (RNS) are often reactive and short-lived, may disrupt the redox state of biological tissues/cells (*e.g.*, red blood cells (RBCs), plasma). Several methods have been developed to detect the redox properties of the blood using the optical^13, 15^ or magnetic properties^16, 17^ of the inorganic iron-chelate of hemoglobin (Hb) and plasma albumin.

Electron spin resonance is commonly used to detect the ROS/RNS directly^18, 19^. However, the approach is hampered by inherent sample stability issues and limited sensitivity^20^. Stable molecular products formed from reactions with ROS/RNS, such as the oxidation targets (*e.g.*, lipid, protein, nucleic acid) are measurable using a range of spectrophometric assays and mass spectrometry (MS)^21^. Nevertheless, fluorescent-staining often causes cell-toxicity^21, 22^, and therefore these assays may not provide information that reflects *in vivo* conditions. Ultraviolet-visible light spectroscopy has poor spectral resolution, and limited sensitivity. Furthermore, globin-associated free radical in Hb is not optically visible^23^ (Supplementary Figures 1-3). MS-based analysis of ROS/RNS reaction products is a powerful and sensitive technique to reveal detailed chemistry of these species, yet requires substantial sample preparation and therefore difficult to be deployed as a rapid screening tool^24^.

We herein report an approach to rapidly quantify the composite redox state of the Hb/plasma by direct measurement of proton relaxation rates of (predominantly) bulk water using a bench-top sized micro magnetic resonance relaxometry (micro MR) system (Figures 1A-B)^25, 26^. The non-destructive nature of the micro MR analysis allows oxidative stress to be artificially introduced in *ex vivo* environment using different biochemical compounds (*e.g.*, nitrite, peroxide) in a controlled manner (Figure 1C). This allows functional assessment of the oxidative susceptibility, tolerance and capacity of a given sample. This yields significantly richer and clinically useful information about the oxidative stress levels of the blood within an individual, as compared to routine biomarkers. To enumerate the various redox states of the Hb (*e.g.*, Fe^2+^, Fe^3+^, Fe^4+^, and globin-associated radical Fe^4+^) and the plasma, two dimensional relaxations map, known as T_1_-T_2_ magnetic state diagram was proposed (Figure 1D). This magnetic state diagram allows visualization and identification of the intermediate redox states and the transient, dynamic pathways of the blood sample.

**Figure 1:**
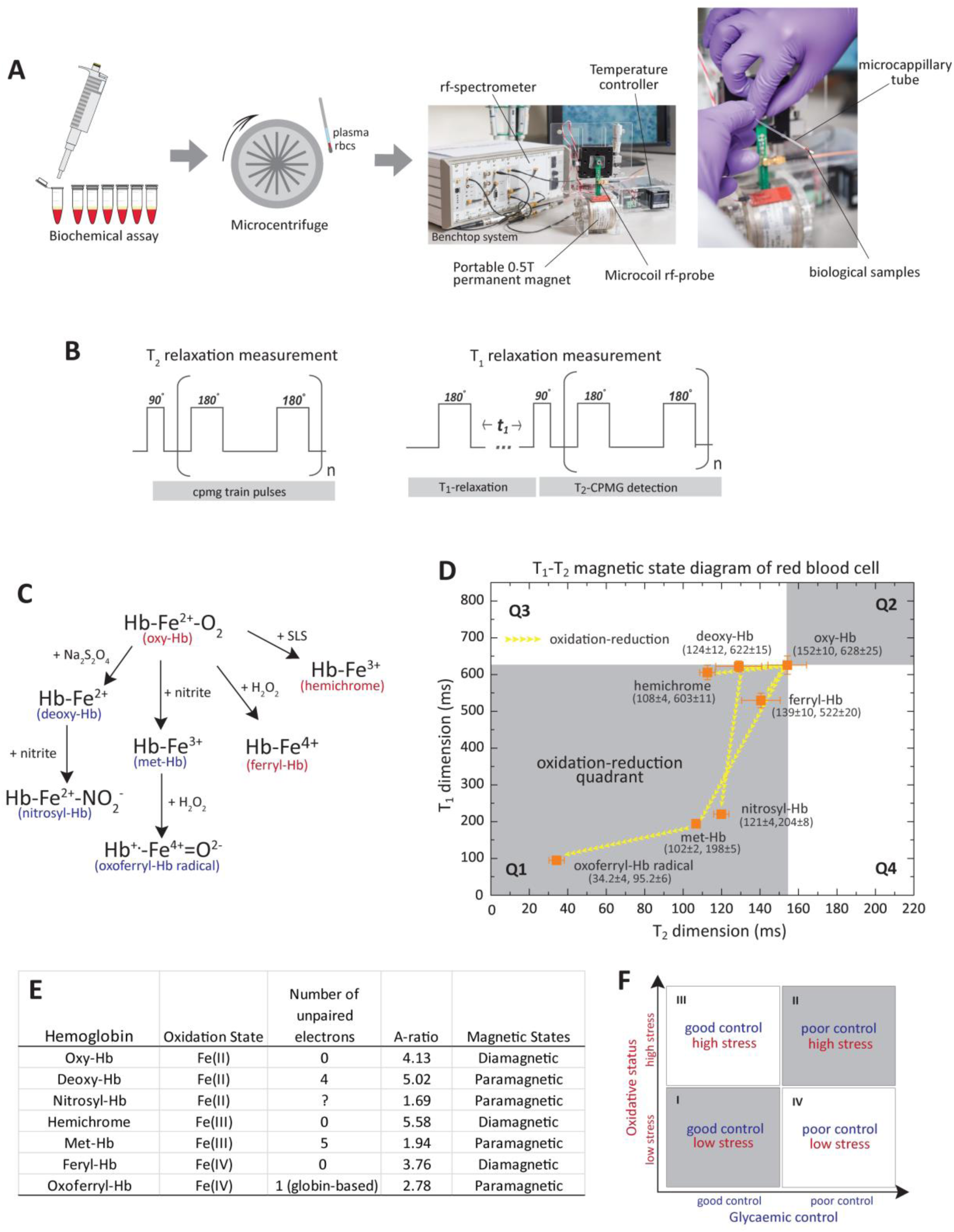
Functional sub-Phenotyping of Oxidative Stress with micro MR analysis approach. (a) Schematic illustration of the micro MR assays performed in this work. Once the patient’s blood is collected via venipuncture, necessary biochemical assay is performed in blood aliquots. Chemical reagent (*e.g.*, nitrite, peroxide) is mixed with the fresh blood and incubated for an interval of 10 min (unless mentioned otherwise) in selected concentration. Micro capillary tubes were then used to sample the biological samples *i.e.*, RBCs/plasma. Standard centrifugal force (3000 *g*, 1 minute) was used to separate and concentrate the packed RBCs from the buffer to avoid possible hematocrit variation in patients. The capillary tubes were then slotted into the rf-probe for micro MR analysis and the read-out completes in less than 5 minutes. Proton NMR of predominantly the bulk water of red blood cells (and plasma) were adjusted to resonance frequency of 21.57 MHz. The portable micro MR system developed in this work consists of a benchtop console, detection circuit coil mounted on a micro stage and a palm-sized 0.5 T permanent magnet, and a temperature controller to stabilize the magnetic field within the chamber and biological sample under measurement. (b) The rf pulse sequences used were standard CPMG pulse sequence and standard inversion recovery experiment (with CPMG detection) for the T_2_ relaxations, and T_1_ relaxations measurements, respectively. In order to obtain high signal-to-noise ratio under relatively inhomogeneous magnetic environment, an array of echoes (a few thousands) within a very short echo interval (in the order of μs) were used to acquire spin-echoes from less than 4 μL sample volume of packed RBCs or plasma. (c) Redox reaction of the iron-heme in various oxidation states: Fe^2+^, Fe^3+^, Fe^4+^ and globin-radical Fe^4+^, which were chemically-induced in *in-vitro* environment (Methods Online). The haemoglobins were in two-possible magnetic states: diamagnetic (red) and paramagnetic state (blue). (d) Various redox states of hemoglobin mapped out using the proposed T_1_—T_2_ magnetic state diagram. The coordinates (in ms) were oxy-Hb (T_2_=152±10, T_1_=628±25), deoxy-Hb (T_2_=124±12, T_1_=622±15), met-Hb (T_2_=102±2, T_1_=198±5), ferryl-Hb (T_2_=139±10, T_1_=522±20), oxoferryl-Hb (T_2_=34.2±4, T_1_=95.2±6), nitrosyl-Hb (T_2_=121±4, T_1_=204±8), and hemichrome (T_2_=108±4, T_1_=603±11). (e) The corresponding magnetic states, number of (un)paired electron, A—ratio, and oxidation state of iron-heme. Three different samplings were taken from the same donor, and the results were reported as mean ± standard error measurement. (f) A quadrant chart of diabetic subject stratified into subgroups based on their oxidative status in association with their average glycaemic levels (*e.g.*, HbA_1c_).

MR relaxometry is a technique to measure relaxation rate, which can be obtained by acquiring spin-echoes of (predominantly) water content of the cells/tissues using conventional nuclear magnetic resonance (NMR) spectroscopy and magnetic resonance imaging (MRI) system. Recent advances in NMR system miniaturization have raised the prospect of applying these techniques in point-of-care diagnostic setting. These applications include immuno-magnetic labeling based detection (*e.g.*, tumor cells^27–29^, tuberculosis^30^ and magneto-DNA detection of bacteria^31^) and label-free micro MR detection of various diseases (*e.g.*, oxygenation^32^/oxidation^26^ level of the blood and malaria screening^25, 33^).

We applied micro MR analysis on whole blood samples to stratify diabetic subjects into subgroups based on their oxidative status levels in association with their glycemic control (Figure 1F). Assessment of oxidative status by measuring the redox state of whole blood was shown to be highly time-and patient specific, revealing information that is potentially critical for clinical diagnostic, monitoring and prognostic purposes.

## RESULTS

### T_1_-T_2_ Magnetic State Diagram

Redox homeostasis is a fundamental biological process, which maintains the balance between ambient anti-oxidant and pro-oxidant activities. The red blood cell is an important biological agent in ameliorating oxidative stress^34, 35^. On the other hand, free heme is one of the major source of redox-active iron, which causes downstream deleterious effect on DNA/protein and RBCs themselves. The fundamental process of oxidative (and nitrosative) stress involve the process of electron transfer, which lead to the eventual formation of oxidized products. The oxidized product is much more stable and measurable using proton NMR relaxometry.

Here, we chemically induced (Figure 1C and Methods Online) and characterized various redox states of the red blood cell and represented them using T_1_-T_2_ magnetic resonance relaxation state diagram (Figure 1D). Each Hb species has specific oxidation states (*e.g.*, Fe^2+^, Fe^3+^, Fe^4+^, globin-associated radical of Fe^4+^ or its’ corresponding complexes) that are bound to specific neighboring proteins, and dissipate energy via unique relaxations mechanism in both the longitudinal (T_1_) and transverse (T_2_) relaxation frames. The T_2_ and T_1_ relaxation times measurement were performed using the standard Carr-Purcell-Meiboom-Gill (CPMG) pulse sequence^36^ and inversion-recovery observed by CPMG, respectively. The pairing of both relaxation times forms a specific T_1_─T_2_ relaxometry coordinate, which is unique to each redox state.

These relaxation times reflect predominantly the bulk water, which came into contact with macromolecular proton (*e.g.*, hemoglobin, albumin)^37^. Water is an attractive natural molecular network probing system as it forms hydrogen bonds with practically all others macromolecule (*e.g.*, protein, metabolite) that are present in human circulation^37, 38^. Therefore, a subtle change of the molecular environment can induce a measurable change in the proton relaxation rate. Among the early works on relaxation rate dependent on the blood oxygenation level were carried out by Thulborn et. al.,^39^ Gomori et. al.,^40^ and eventually used to measure brain activity known as functional MRI^41^.

Oxyhemoglobin (oxy-Hb) which has the lowest reduced ferrous (Fe^2+^) state is the predominant Hb species in circulation. The oxy-Hb can be provisionally assigned to the center of the state diagram, which has four quadrants (*i.e.*, Q1, Q2, Q3 and Q4). Due to the semi-solid structure of RBC and oxidation process which reduces the proton relaxation time, the redox pathways of RBC mapped out predominantly in Q1.

Electrons in the *d* sub-orbital of iron hemoglobin can exist in various paired or unpaired conditions, rendering them into two possible magnetic states, *i.e.*, diamagnetic and paramagnetic states, respectively. Hb with at least one unpaired electron, *i.e.* deoxygenated hemoglobin (deoxy-Hb), methemoglobin (met-Hb), nitrosyl hemoglobin (nitrosyl-Hb), and oxo-ferryl radical exhibit the effect of paramagnetism with much larger bulk magnetic susceptibility than its’ diamagnetic counterparts *i.e.*, oxy-Hb, ferryl-Hb, and hemichrome (HC) (Figures. 1D-E, Methods Online). The magnetic relaxivity contributed by paramagnetic ion is highly dependent on its spin state, and is directly proportional to *S(S+1)*, where *S* is the spin quantum number of the total electron spin. Each of the Hb oxidation states has a specific normalized relaxation constant (A-ratio=T_1_/T_2_) which represent a unique identifier (Figure 1E).

### Nitrite-induced Ferrous Oxidation

Freshly collected whole blood samples containing predominantly the oxygenated Hb were oxidized *in-vitro* into met-Hb in the presence of sodium nitrite (Methods Online). The oxygenation levels generated *in vitro* were independently verified using spectrophotometry (Supplementary Figure 1A). Redox titration profile showed a strong dose-dependent curve, where both T_1_ and T_2_ relaxation times reduced gradually as progressively higher proportion of RBCs were oxidized and increased the volume paramagnetic susceptibility, when the nitrite concentrations were increased from 50 nM to 10 mM (Figures 2A-B). As the blood sample transformed to a complete paramagnetic state (T_2_=92.8 ms, T_1_=190.0 ms) from the initial diamagnetic states (T_2_=149.0 ms, T_1_=620.0 ms), the A-ratio dropped from 4.16 to 2.02 (R^2^>0.95, Figure 2C). As the volume paramagnetic susceptibility increased, this causes the T_1_-T_2_ trajectory to move downward in Q1 (Figure 2D).

**Figure 2:**
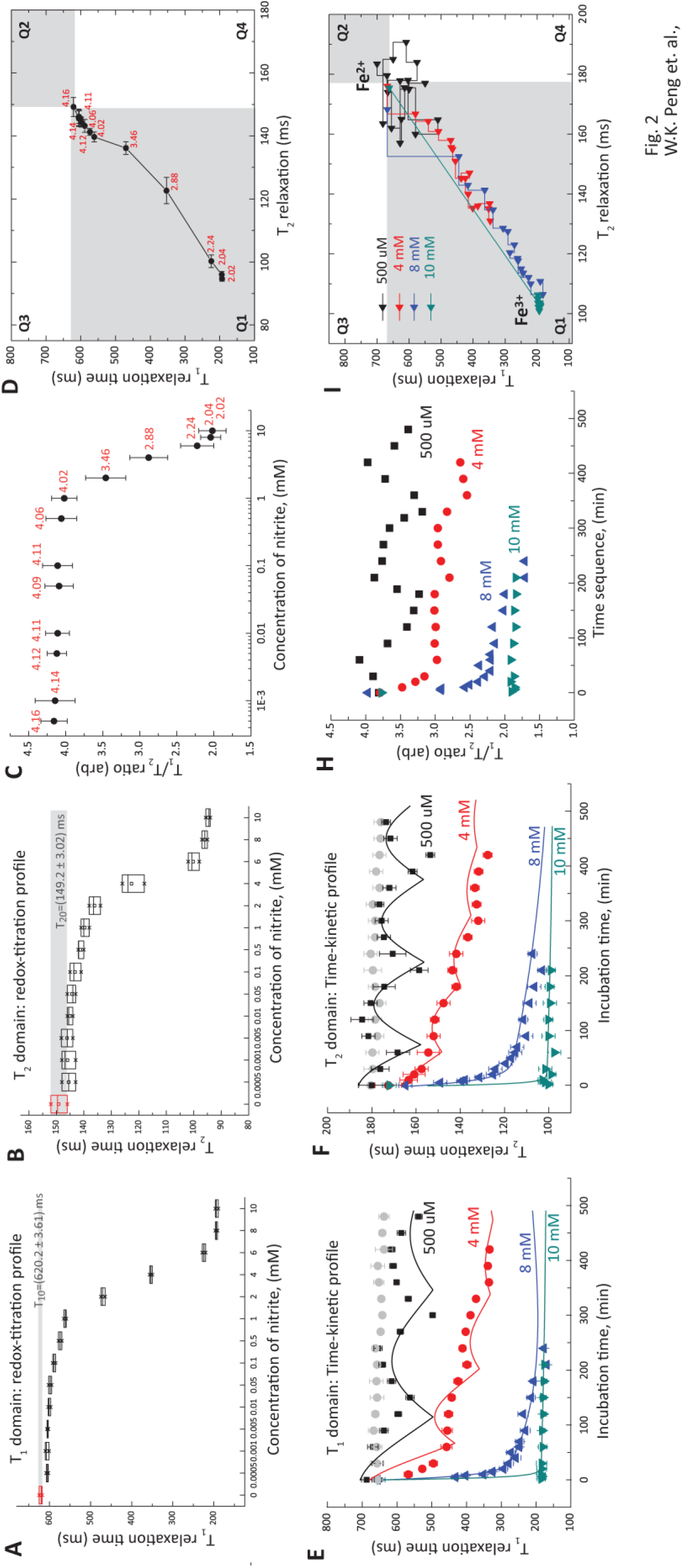
Nitrite induced ferrous oxidation: Redox-titration profile of red blood cells as function of nitrite concentration in (a) T_1_ relaxation and (b) T_2_ relaxation domain. The incubation times were 10 minutes. The control baseline readings were (T_2o_=149.5, T_1o_=621.3) ms, which is the readings for oxy-Hb without any nitrite exposure. The corresponding concentration dependent (c) A—ratio, and (d) T_1_—T_2_ trajectories of the gradual inversion of Fe^2+^ subpopulation to complete formation of Fe^3+^ population. Time dependent kinetic profile of ferrous oxidation using nitrite concentrations (500 μM, 4 mM, 8 mM and 10 mM) in (e) T_1_ relaxation and (f) T_2_ relaxation domain. The corresponding (c) A— ratio, and (d) T_1_—T_2_ trajectories in the magnetic state diagram. Three different samplings were taken from the same donor, and the results were reported as mean ± standard error measurement.

The dose-dependent reaction was lost when excess of nitrite (>10 mM) was introduced. This suggests that the oxidant concentration had exceeded anti-oxidant capacity and all the possible oxidation states were saturated. At much lower concentration (<100 µM), there was little or no change in the bulk magnetic state of the RBCs, as the majority of the RBCs were able to restore to their original reduced state. Interestingly, steep transitional oxidation zone was observed within a very narrow range of nitrite concentration; from 1 mM to 8 mM, which reflected the redox homeostatic responses within the concentration where the cells were viable. This was crucial to the understanding of the functioning of RBCs at cellular and subpopulation levels (Figures 2A—C).

Further evidence of redox homeostasis was observed in time-dependent kinetic profiles (Figures 2E—F) over a range of nitrite concentrations (500 µM, 4 mM, 8 mM and 10 mM). In general, the measured T_1_ and T_2_ readings changed in an oscillatory manner over time. This may suggest an active mechanism to regulate cell redox homeostasis. As the RBCs aged, antioxidant capacity is reduced, thereby forming a subpopulation of cell with disproportionately low antioxidant capacity^1^.

The amplitudes of the oscillation decreased as the nitrite concentration was increased from 500 µM to 4 mM (Figure 2H). At much higher nitrite concentration (>10 mM), the reaction curve decayed rapidly in an exponential manner with an increasingly dampened oscillation. Similar observations were recorded using spectrophotometry (Supplementary Figure 1).

Interestingly, the corresponding kinetic profiles followed an identical path over time in the T_1_-T_2_ trajectories as the nitrite concentration was increased (Figure 2G). The oxidation process drove all the trajectories toward a common coordinate (T_2_= 92.8 ms, T_1_= 190.0 ms), where all the RBCs were converted fully into met-Hb. For low nitrite concentration (*e.g.*, 500 μM) however, the T_1_-T_2_ trajectory circulated around the origin and did not reach the eventual met-Hb coordinates.

### Functional Phenotyping of Oxidative and Nitrosative Stress in Subjects with Diabetes Mellitus

A cross sectional study was carried out to stratify DM subjects based on their oxidative status. DM subjects (n=185) who had HbA_1c_ measured in the outpatient clinic as part of their clinical care (random blood sample) were included in this study. These subjects had HbA_1c_ ranging from 4% to 16% and the subjects were classified into good glycaemic control (<7.0% HbA_1c_) and poor glycaemic control (>8.0% HbA_1c_) subgroups^2^. Healthy young male subjects (n=32; age range of 21 to 40 years, fasting glucose below 5.6 mmol/L, average HbA_1c_ of 5.16 (±0.32) %, and body mass index below 23.5 kg/m^2^) were separately recruited as control subjects. The collected whole blood in EDTA-anticoagulated tubes were centrifuged (14 000 *g*, 5 min) to separate the RBCs and plasma. The micro MR analysis was performed blindly on freshly collected fasting blood samples or otherwise kept at 4°C within 2 hours. Other clinical laboratory tests (*e.g.*, HbA_1c_) were performed in parallel.

### Baseline study: Oxidative Status of Glycated Hb in RBCs

The baseline reading of intact RBCs were measured and mapped using in T_1_-T_2_ magnetic state diagram. It appears that subjects with poor glycaemic control (red) have much shorter T_1_ and T_2_ readings as compared healthy control subjects (blue) (Figures 3A-B). This was mainly due to the presence of higher concentration of ferric Hb *i.e.*, the low spin (HC) and high spin (met-Hb), which contribute to reduction in T_1_ and T_2_ relaxation times (Figure 2 and Supplementary Figure 4). The formation of HC and its relaxation response were further verified using *in-vitro* chemical stress (Supplementary Figure 4). In particularly, ferric Hb concentration ware markedly elevated in DM subjects with poor glycaemic control (>10% HbA_1c_) (Figures 3B—D). Although DM subjects with good glycaemic control had only slightly elevated T_1_ and T_2_ readings, as compared with healthy control subjects (Figure 3b), the use of A-ratio plot yielded significantly better resolution (*P*<0.01) (Figure 3E).

**Figure 3:**
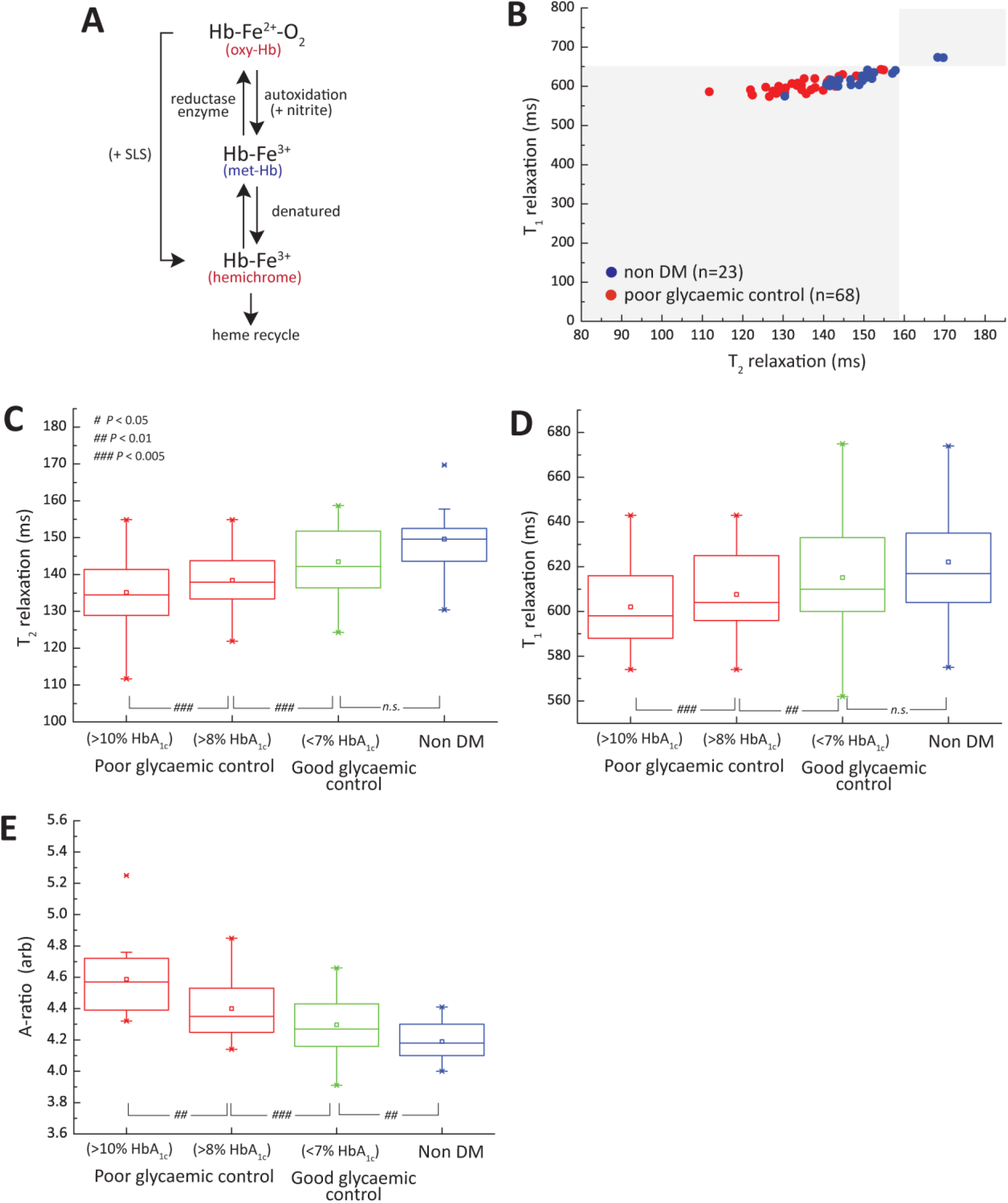
Functional Phenotyping of Nitrosative Stress in RBCs for Subjects with Diabetes Mellitus. (a) *In vivo* **r**edox formation of high-spin met-Hb, low-spin hemichrome and heme metabolism. The equivalent chemically induced *in-vitro* environment using sodium salicylate (SLS). (b) The T_1_—T_2_ relaxometry coordinates of RBCs baseline readings of non-DM subjects (blue, n=23) and subjects with poor glycaemic control (red, n=68). (c) T_2_ relaxation and (d) T_1_ relaxation domain and the corresponding (e) A-ratio index of subjects with poor glycaemic controls (n=62) and good glycaemic control (n=50) subgroup as compared to healthy non DM subjects (n=20). The subjects with poor glycaemic controls were further sub-divided into >8% HbA_1c_ (n=47) and >10% HbA_1c_ (n=15) subgroups. The statistical significance was calculated using the Student’s T-Test (two-tailed, unequal variance).

**Figure 4:**
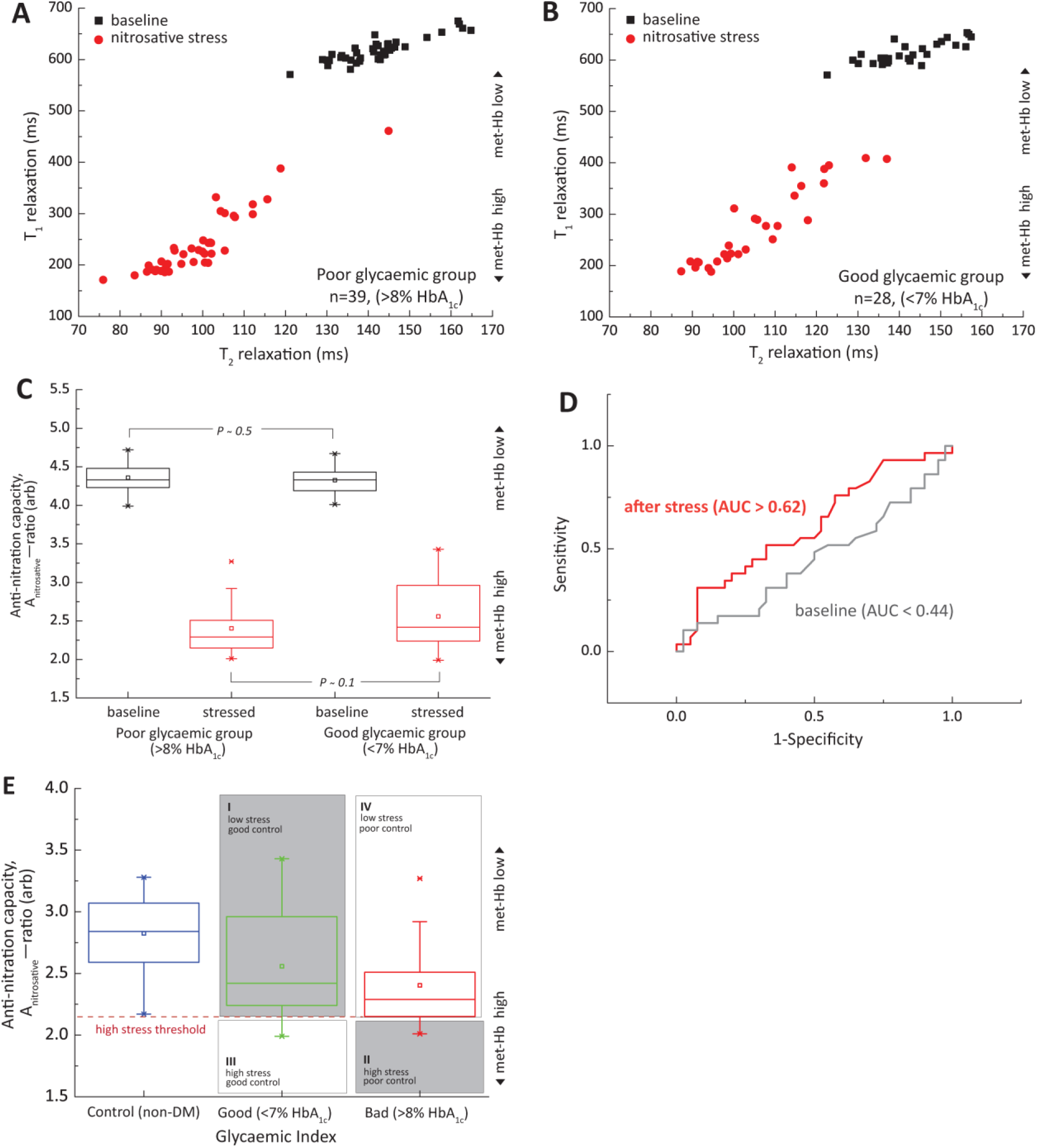
*Ex vivo* Nitrosative Functional Stress Test on Glycated-Hb. The T_1_—T_2_ relaxometry coordinates of RBCs taken before (black) and after (red) nitrite treatment for subjects with (a) poor glycaemic control (n=39), and (b) good glycaemic control (n=28). (c) Its’ corresponding distribution based on A—ratio index. The statistical significance was calculated using the Student’s T-Test (two-tailed, unequal variance). (d) The diagnostic accuracy as calculated using ROC curve for RBCs taken before (black) and after (red) the stress test. The probability diagnostic accuracy is quantified as Area Under the Curve (AUC). (e) A quadrant chart of diabetic subjects stratified into subgroups based on their oxidative status (nitrosative stress) in association with their glycemic levels (*e.g.*, HbA_1c_) as compared with healthy non DM subjects (n=23). The proposed method segregated effectively the subgroup III subjects (good glycemic control and yet high nitrosative stress) from the rest of the cohorts. Note that the Y-axis (nitrosative stress) were inversely represented as compared to quadrants shown in Figure 1F.

### *Ex vivo* Nitrosative Functional Stress Test on Glycated-Hb

To further evaluate the ability of RBCs to tolerate the nitrosative stress, we artificially challenged the RBCs with strong oxidant (nitrite). Freshly collected RBCs were incubated with 6 mM sodium nitrite for 10 min, washed three times to stop the reaction and finally resuspended in *1x* PBS for micro MR analysis (Methods Online). The sodium nitrite concentration chosen was within the viable homeostatic range (Supplementary Figures 5, 6 and 7). The micro MR analyses were assessed before (black square) and after (red circle) the stress test.

**Figure 5:**
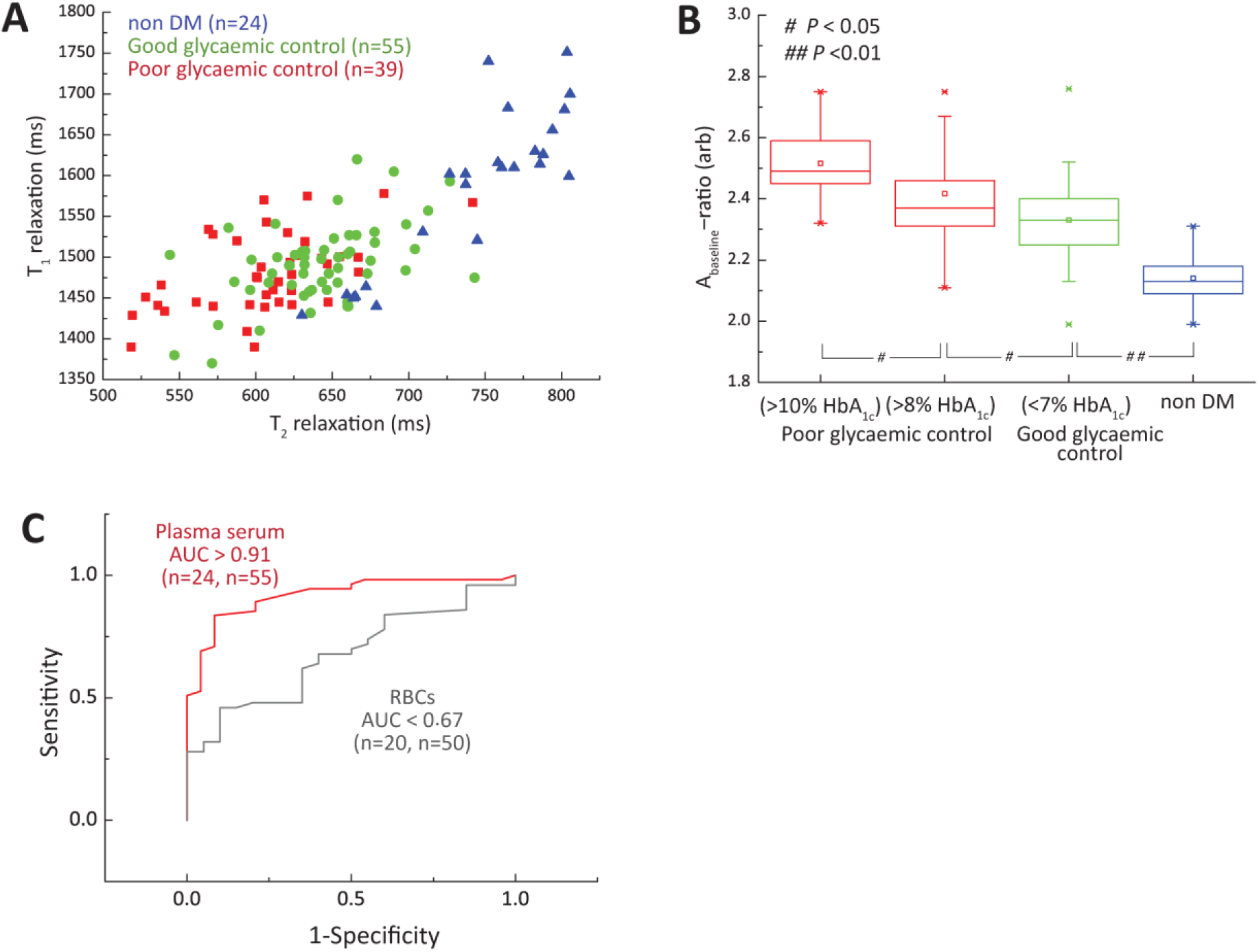
Functional Phenotyping of Oxidative Stress in Plasma for Subjects with Diabetes Mellitus. (a) The T_1_—T_2_ relaxometry coordinates of plasma baseline taken from healthy non DM subjects (blue, n=24), subjects with good glycaemic control (green, n=55) and subjects with poor glycaemic control (red, n=39). (b) The corresponding A-ratio against the subjects with poor glycaemic control (n=39) and good glycaemic control (n=55) subgroups, as compared to healthy non DM subjects (n=24). The subjects with poor glycaemic controls were further subdivided into >8% HbA_1c_ (n=14) and >10% HbA_1c_ (n=25) subgroups. The statistical significance was calculated using the Student’s T-Test (two-tailed, unequal variance). (c) The diagnostic accuracy of RBCs (gray) and plasma (red) taken from subjects with good glycaemic control with respect to healthy non DM subjects. The number of subjects (n) were indicated on the parentheses (non-DM, good glycaemic control).

**Figure 6:**
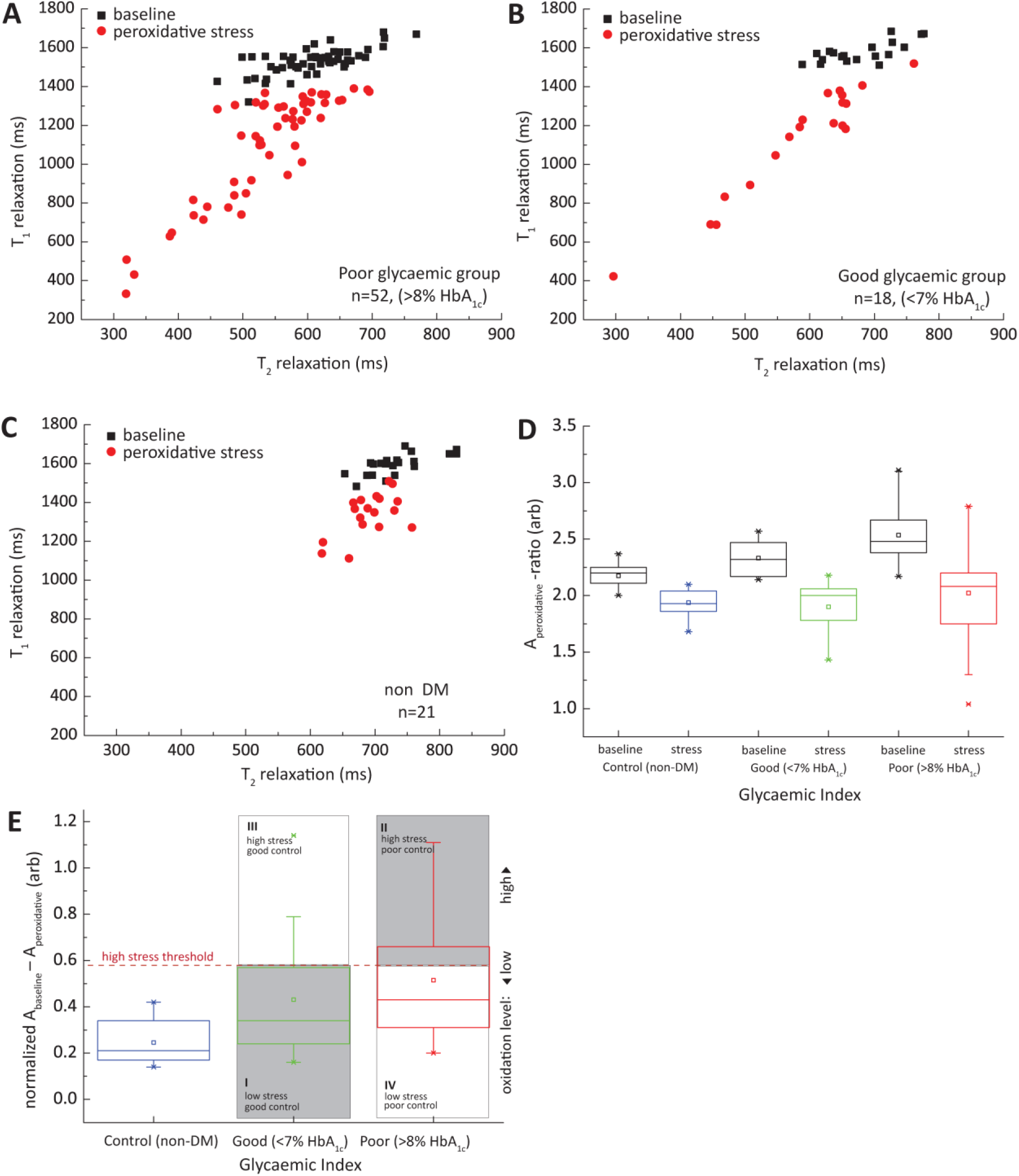
*Ex vivo* Peroxidative Functional Stress Test on Plasma. The T_1_—T_2_ relaxometry coordinates of plasma baseline (black square) and plasma pretreated with hydrogen peroxide (red dot) for subjects with (a) poor glycaemic control (n=52), (b) good glycaemic control (n=18), and (c) healthy non DM (n=21). (d) The corresponding A_peroxidative_ index taken before (black) and after (blue, green, red) the peroxidative stress test. (e) A quadrant chart of diabetic subjects stratified into subgroups based on their peroxidative status (normalized A_baseline_ — A_stress_) in association with their glycaemic levels (*e.g.*, HbA_1c_) as compared with healthy non DM subjects. Note that the Y-axis (peroxidative stress) were inversely represented as compared to Y-axis (nitrosative stress) in quadrant shown in Figure 4E.

DM subjects with poor (n=39) and good (n=28) glycaemic control with similar matching RBCs baseline (black squares in Figure 4A-B) were chosen for this test. Lower anti-nitration capacity (or increased in nitrosative stress susceptibility) was indicated by an increase in met-Hb formation (red circles) and hence the lower the A_nitrosative_-ratio value. DM subjects vary markedly in their nitrosative susceptibility despite having similar baseline; with DM subjects with poor glycaemic control being more susceptible to nitrite-induced oxidation as compared to its’ counterpart with good glycaemic control (Figure 4C). Receiver Operating Characteristic (ROC) analyses showed that the initial baseline (AUC<0.44) between the subjects from poor and good glycaemic subgroups had increased (AUC>0.62) upon nitrite stress (Figure 4D). Due to structural modification of Hb as a result of increased glycation, HbA_1c_ is less stable and more prone to oxidation, in agreement with observation reported elsewhere^42, 43^.

The spread of the baseline was large for the good glycaemic control group, which suggests a large between-subject variability of nitrosative susceptibility, despite having similar glycaemic level (Figure 4E). Using the nitrosative susceptibility (A_nitrosative_-ratio), which could be derived hypothetically from this study and the traditional index of glycaemic control (HbA_1c_), DM subjects could be stratified into four distinct quadrants (*i.e.*, Q1 to Q4). This approach singled out a minority group in Q3 (subgroup III), who had good glycaemic control, and yet had (high) nitrosative stress A_nitrosative_-ratio that was at 75^th^ percentile that of typical DM subjects with poor glycaemic control and at 95^th^ percentile of the healthy control subjects.

### Baseline Study: Glycation and Glycoxidation of Plasma

Increased blood glucose promotes non-enzymatic glycation of plasma proteins, which include the albumin, alpha-crystalline, collagen, and low-density lipoprotein. A large proportion of total serum protein is attributable to serum albumin^44, 45^. Glycation and oxidative damage cause protein modification, which affects the protein functionality^46^. The micro MR analyses were performed at room temperature (26°C). Each T_1_-T_2_ coordinate represents the composite redox properties of one subject’s plasma (Figure 5A). The baseline readings of the DM subjects have much shorter T_1_ and T_2_ relaxation times, and it was well separated from the healthy non DM subjects (blue). Notably, DM subjects with poor glycemic control, in particularly DM subjects of >10% HbA_1c_ subgroup (mean A-ratio of 2.52), seen a strong departure from the healthy controls (mean A-ratio of 2.13) (Figure 5B).

The marked reduction in relaxation states was attributed to an increase in glycation and glycoxidation of the serum albumin, known as glucose toxicity. As a result of increased glycation (*in-vitro* validation in Supplementary Figure 8a) and protein oxidative damage^46^ (*e.g.*, protein cross-linking), the bulk water proton of mobility was further restricted,^47^ leading to reduction in T_1_ and T_2_ relaxation times. T_2_ relaxation however reduced much faster than T_1_ relaxation resulting in an increase in A_baseline_-ratio, which correlated positively with HbA_1c_ (R^2^>0.2) (results not shown). Similar trends were observed *in vitro*, which confirm the effects of glycation (Supplementary Figures 8A-B) and glycoxidation (Supplementary Figure 8C). A separate study by Cistola *et. al.*, recently found that the baseline T_2_ of water plasma/serum shown a strong correlation in subjects with metabolic abnormalities^38, 48^. Interestingly, ROC analysis indicates DM subjects with good-glycemic control and healthy controls subjects had AUC of 0.91, much higher than the one observed in RBCs (AUC<0.67) (Figure 5C). The result seems to suggest that pathological footprint of hyperglycemia is more prominent in extracellular plasma as compared to the RBCs at baseline level.

### Peroxide induced Oxidation Analysis: Total Anti-oxidant Capacity of Plasma in Diabetes Mellitus Subjects

In order to evaluate the total anti-oxidant capacity of plasma towards oxidation, we artificially challenged the plasma with hydrogen peroxide in subjects with poor glycaemic control (n=52), good glycaemic control (n=18), and healthy control (n=21) (Figure 6). Hydrogen peroxide solution was added into the freshly drawn plasma (10% v/v) for an incubation time of 10 min (Methods Online and Supplementary Figure 9). The micro MR analyses were performed before (black squares) and after (red circles) the mixing (Figures 6A—C).

The results of this stress test revealed a large spread of T_1_-T_2_ coordinates for DM subjects (Figures 6A—B), indicating marked variation in their peroxidative susceptibility as compared to healthy controls (Figure 6C). Lower anti-oxidant capacity (or increase in peroxidative stress susceptibility) of plasma is indicated by reduction in T_1_ and T_2_ relaxation coordinates (red circles). As more oxidized plasma was formed, the T_1_ relaxation time reduced much faster than T_2_ relaxation time, and hence the reduction in A_peroxidative_-ratio (Figure 6D), which was in agreement with *in vitro* validation (Supplementary Figure 8c).

DM subjects had much higher plasma peroxidative susceptibility as compared to non-DM counterparts (Figure 6E). The normalized plasma peroxidative stress susceptibility can be defined by the difference between the A_baseline_-ratio and the A_peroxidative_-ratio (Figure 6e). Note that the plasma baseline (black) measurements of this cohort having similar positive correlation with glycaemic levels (Figure 6D) were in agreement with the previous cohort measured independently^49^ (Figure 5). Exposure to peroxyl compound leads to an increased formation of disulfide bonds in albumin and human non-mercaptalbumin, which was also observed in several others pathological state^50, 51^. The proposed peroxidative susceptibility measurement that is independent of HbA_1c_ can be used to stratify the DM subjects into subgroups, which provide insight into the oxidative status (susceptibility and damage) in personalized manner (Figure 6E).

## DISCUSSION

We have developed a highly sensitive approach to accurately detect and quantify the redox (and hence oxidative/nitrosative) state and the subtle molecular motion changes of blood samples, inferred based on the relaxation measurement. This is the first demonstration of the unique magnetic resonance relaxation properties of the various hemoglobin states, which were mapped out using the proposed magnetic state diagram. The measurement of redox properties in plasma/erythrocytes can provide a useful parameter for functional phenotyping of many biological pathways to better understand disease pathophysiology. This technology has vast potential to be applied for clinical disease diagnosis, prognosis and monitoring, given that the specificity of the oxidative stress in association with the disease state can be further improved in near future.

The platform presented here has several innovative features and is readily adaptable for clinical use (Supplementary Figure 10). Firstly, the miniaturized platform^26, 52^ developed here is portable and the proposed assays requires minimal processing steps, low-cost, robust and can therefore be performed by minimally trained operator. The high sensitivity can be attributed to the micron-sized detection coil and optimized ultra-short echo time implemented in this work. Only a minute amount of blood sample volume (< 10 µL) is needed for each test, which enables the collection of sample using minimally invasive technique^26, 53^ such as finger prick a standard procedure in patient care.

Secondly, we exploited the non-destructive nature of magnetic resonance, and introduced a number of *in-vitro* functional assays that yielded parameters about the oxidative status of an individual, which may be clinically useful. It probes the primary redox event as compared to the current gold-standard biomarker, isoprostanes, which is a downstream marker and may be susceptible to confounding factors. The use of isoprostanes as biomarker of oxidative status for correlation with disease outcome has so far yielded conflicting results in cross-sectional versus longitudinal studies^54, 55^. Furthermore, they are static biomarkers that provide snapshots of the oxidative status of biological samples representing the *in vivo* condition of the subject at the point of collection. To accurately measure these molecules, laborious technique such as gas-or liquid-chromatography mass spectrometry has to be employed, limiting its’ utility as diagnostic tools.

Further clinical validation is needed to compare current proposed biomarkers with isoprostanes, and a combined assessment may yield even richer information. A long-term follow-up and large-scale prospective study is currently underway to evaluate the diagnostic performance of this technique. This accurate and rapid technique for quantification of oxidative stress may be included in future risk stratification models where subjects with single or multiple complications can be streamlined based on their oxidative index. This work opens up new opportunities for molecular phenotyping of oxidative stress in a rapidly and systematic manner for various chronic diseases (*e.g.*, cancer) and a range of hematology applications (*e.g.*, sepsis), including the acquired and congenital diseases such as enzymatic deficiency, Hb synthesis defects (*e.g.*, Thalassemia), and Hb molecular defects (*e.g.*, sickle cells anemia, unstable Hb).

## ACKNOWLEDGEMENT

This research was supported by Start-Up Grant of W.K. Peng at International Iberian Nanotechnology Laboratory and National Research Foundation Singapore through the Singapore MIT Alliance for Research and Technology’s BioSystems and Micromechanics Inter-Disciplinary Research programme. BOB is supported by grants from LKC School of Medicine.

## AUTHOR CONTRIBUTIONS

W.K. Peng and T.P. Loh conceived the original idea and analyzed the results, and wrote the first draft of the paper together. W.K. Peng designed the experiments/protocols, proposed the magnetic state diagram, and spearhead the entire hardware development. L. Chen assists in hardware development and performed most of the micro MR analyses and related assays. B.O.B provided input regarding translational medicine applications. All the authors checked through the manuscript and analyzed the data.

## METHODS and MATERIALS

### Magnetic resonance relaxation measurement and detection

The proton nuclear magnetic resonance (NMR) measurements of predominantly the bulk water of red blood cells (RBCs) and plasma were carried out at the resonance frequency of 21.57 MHz using a bench-top type console (Kea Magritek, New Zealand) and portable permanent magnet (Metrolab Instruments, Switzerland), B_o_=0.5 T. A homebuilt temperature controller was constructed to regulate the temperature at 26°C inside the measurement chamber. This helped maintained the stability of the magnetic field and biological sample under measurement. Single resonance proton micro magnetic resonance (MR) probe with detection coil of 1.55 mm inner diameter was constructed to accommodate a heparinized microcapillary tube (outer diameter: 1.50 mm, inner diameter: 0.95 mm) (Fisherbrand, Fisher Scientific, PA) for a detection region of approximately 3.8 µL in volume. The longitudinal relaxation times, T_1_ were measured by standard Inversion-Recovery pulse sequences observed by Carr-Purcell-Meiboom-Gill (CPMG) train pulses. The transverse relaxation times, T_2_ were measured by standard CPMG train pulses (inter echo time: 200 µs) consisting of 2000 echoes. A total of 12 scans were typically acquired for signal averaging unless mentioned otherwise. The transmitter power output was maintained at 360 mW for a single 90° pulse of 6 µs pulse length, which corresponds to a nutation frequency of 41.6 kHz. The delay between each pulse (recycle delay) were set at 1s and 4s for RBCs and plasma, respectively.

### Ethics and Blood Collection

This study received approval from the local Institutional Review Board of the National Healthcare Group. Patients were not identified throughout the study. The EDTA-anticoagulated whole blood samples were collected using standard phlebotomy procedures. All blood samples were kept at ≤4**°**C within two hours of collection and were kept refrigerated until analysis.

### Healthy subjects

Subjects without past history of diabetes mellitus (DM) and had normal oral glucose tolerance test according to the American Diabetes Association criteria (fasting glucose <5.6 mmol/L; two hour post oral glucose tolerance test glucose of <7.8 mmol/L) were recruited into this study following provision of informed consent. They were Chinese males aged between 21 and 40 years, with a body mass index below 23.5 kg/m^2^.

### Subjects with Diabetes Mellitus

Annonymised residual samples collected from DM patients at the outpatient clinic for measurement of glycated hemoglobin (HbA_1c_) as part of their clinical care, were included in this study. The HbA_1c_ was measured using the Bio-Rad Variant II analyzer. This National Glycohemoglobin Standardization Program (NGSP) certified instrument has an analytical coefficient of variation of <2% at HbA_1c_ concentration of 4% and 16%. Our laboratory is NGSP level 1-certified.

### Sample Preparation and micro MR analysis

Fresh RBCs were washed three times with 1*x* PBS solution and re-suspended at 10% hematocrit with PBS. The selected chemical was then mixed into the prepared blood at desired concentration (see **Biochemical Assays** details below). The final concentrations were recalculated based on the entire volume. Other lower concentrations were prepared according to appropriate dilution. A horizontal shaker was used to homogenously mix (at 200 rpm) for all the chemically treated samples at room temperature. The blood was incubated between a few minutes to a few hours, as indicated in *Text*. The blood was then washed three times to remove the chemical residual. A heparinized microcapillary tube is used to transfer 40 µL volume of blood via capillarity action. In order to obtain packed RBCs for micro MR analysis, the microcapillary tubes were spun down at 3000 *g* for 1 minute.

### Details on Biochemical Assays. Sodium nitrite treated RBCs

20 µL of the desired concentration (in the range 500 µM to 100 mM) of sodium nitrite were then mixed into 180 µL of the prepared blood. Hydrogen peroxide treated RBCs. 20 µL of 3% hydrogen peroxide stock solution (approximately 0.9 M), which was purchased commercially from Sigma Aldrich, was mixed into 180 µL of the prepared blood. Sodium salicylate treated RBCs. 20 µL of the desired concentration (as described in the *Text*) of sodium salicylic were then mixed into 180 µL of prepared blood. Preparation of Oxo-ferryl Hb. Oxo-ferryl Hb was prepared in two steps. The RBCs were first treated with sodium nitrite (similar to the protocol described above) to convert the RBCs into met-Hb. Hydrogen peroxide were then added into the met-Hb using the same protocol as described above. Preparation of Nitrosyl-Hb. The nitrosyl Hb was prepared in two steps. The RBCs were first converted into deoxygenated Hb (similar to the protocol described below) and treated with sodium nitrite using the same protocol as described above. Preparation of Deoxygenated Hb. 20 µL of natrium hydrosulfite, Na_2_S_2_O_4_ (10 mM final concentration, Sigma Aldrich) were then mixed into 180 µL of prepared blood and mix homogenously (at 200 rpm) for 10 minutes with a horizontal shaker. The UV-Visible spectrum was recorded immediately to confirm the presence of deoxygenated hemoglobin. Pure gas N_2_ was continuously purged into an airtight chamber in order to maintain the de-oxygenated condition. The UV-VIS absorbance was used to confirm the presence of deoxygenated-Hb by its distinct peak at 543 nm. Hydrogen peroxide treated plasma. The fresh whole blood collected were centrifuged at 14,000 *g* for 5 minutes to separate the plasma from the packed RBCs. 10 µL of hydrogen peroxide solution were then mixed into 90 µL of prepared plasma and other lower concentrations were prepared according appropriate dilutions.

### Statistical analysis

Unless otherwise noted, all statistical analyses were performed using OriginPro (OriginLab Corporation, United States). For statistical analysis, t-tests were used. All error bars represent were either in standard deviation (s.d.) or standard error measurements (s.e.m) of means and the statistical results were stated as *P*-values.

### Data Sharing Statement

The original data that support the findings of this study are available from the corresponding author upon reasonable request.

## References

1. Abdulbari Bener FM. Projection of Diabetes Burden through 2025 and Contributing Risk Factors of Changing Disease Prevalence: An Emerging Public Health Problem. Journal of Diabetes & Metabolism. 2014;05(02).

2. Use of Glycated Haemoglobin (HbA1c) in the Diagnosis of Diabetes Mellitus: Abbreviated Report of a WHO Consultation. Geneva: World Health Organization; 2011.

3. Duckworth W, Abraira C, Moritz T, et al. Glucose Control and Vascular Complications in Veterans with Type 2 Diabetes. New England Journal of Medicine. 2009;360(2):129–139.

4. Saisho Y. Glycemic variability and oxidative stress: a link between diabetes and cardiovascular disease? Int J Mol Sci. 2014;15(10):18381–18406.

5. Intensive Blood Glucose Control and Vascular Outcomes in Patients with Type 2 Diabetes. New England Journal of Medicine. 2008;358(24):2560–2572.

6. Zhao H-L, Lai FMM, Tong PCY, et al. Prevalence and clinicopathological characteristics of islet amyloid in chinese patients with type 2 diabetes. Diabetes. 2003;52(11):2759–2766.

7. Boehm BO, Schilling S, Rosinger S, et al. Elevated serum levels of N(epsilon)-carboxymethyl-lysine, an advanced glycation end product, are associated with proliferative diabetic retinopathy and macular oedema. Diabetologia. 2004;47(8):1376–1379.

8. Rabbani N, Thornalley PJ. Measurement of methylglyoxal by stable isotopic dilution analysis LC-MS/MS with corroborative prediction in physiological samples. Nature Protocols. 2014;9(8):1969–1979.

9. Bierhaus A, Humpert PM, Morcos M, et al. Understanding RAGE, the receptor for advanced glycation end products. Journal of Molecular Medicine. 2005;83(11):876–886.

10. Soro-Paavonen A, Watson AMD, Li J, et al. Receptor for Advanced Glycation End Products (RAGE) Deficiency Attenuates the Development of Atherosclerosis in Diabetes. Diabetes. 2008;57(9):2461–2469.

11. Baynes JW. Role of oxidative stress in development of complications in diabetes. Diabetes. 1991;40(4):405–412.

12. Maritim AC, Sanders RA, Watkins JB. Diabetes, oxidative stress, and antioxidants: A review. Journal of Biochemical and Molecular Toxicology. 2003;17(1):24–38.

13. Holley AE, Cheeseman KH. Measuring free radical reactions in vivo. Br. Med. Bull. 1993;49(3):494–505.

14. Ihnat MA, Thorpe JE, Ceriello A. Hypothesis: the ‘metabolic memory’, the new challenge of diabetes. Diabetic Medicine. 2007;24(6):582–586.

15. Shah SS, Diakite SAS, Traore K, et al. A novel cytofluorometric assay for the detection and quantification of glucose-6-phosphate dehydrogenase deficiency. Sci Rep. 2012;2:299.

16. Kopáni M, Celec P, Danisovic L, Michalka P, Biró C. Oxidative stress and electron spin resonance. Clin. Chim. Acta. 2006;364(1–2):61–66.

17. Lee M-C-I. Assessment of oxidative stress and antioxidant property using electron spin resonance (ESR) spectroscopy. J Clin Biochem Nutr. 2013;52(1):1–8.

18. Emanuel NM, Saprin AN, Shabalkin VA, Kozlova LE, Krugljakova KE. Detection and Investigation of a New Type of ESR Signal characteristic of Some Tumour Tissues. Nature. 1969;222(5189):165–167.

19. Svistunenko DA, Patel RP, Voloshchenko SV, Wilson MT. The Globin-based Free Radical of Ferryl Hemoglobin Is Detected in Normal Human Blood. Journal of Biological Chemistry. 1997;272(11):7114–7121.

20. Takeshita K, Ozawa T. Recent progress in in vivo ESR spectroscopy. J. Radiat. Res. 2004;45(3):373–384.

21. Buckman JF, Hernández H, Kress GJ, et al. MitoTracker labeling in primary neuronal and astrocytic cultures: influence of mitochondrial membrane potential and oxidants. J. Neurosci. Methods. 2001;104(2):165–176.

22. Spasojević I, Bajić A, Jovanović K, Spasić M, Andjus P. Protective role of fructose in the metabolism of astroglial C6 cells exposed to hydrogen peroxide. Carbohydrate Research. 2009;344(13):1676–1681.

23. Rifkind JM, Abugo O, Levy A, Heim J. [28] Detection, formation, and relevance of hemichromes and hemochromes. Methods in Enzymology. 1994;231:449–480.

24. Aebersold R, Mann M. Mass spectrometry-based proteomics. Nature. 2003;422(6928):198–207.

25. Peng WK, Kong TF, Ng CS, et al. Micromagnetic resonance relaxometry for rapid label-free malaria diagnosis. Nature Medicine. 2014;20(9):1069–1073.

26. Peng WK, Chen L, Han J. Development of miniaturized, portable magnetic resonance relaxometry system for point-of-care medical diagnosis. Review of Scientific Instruments. 2012;

27. Castro CM, Ghazani AA, Chung J, et al. Miniaturized nuclear magnetic resonance platform for detection and profiling of circulating tumor cells. Lab Chip. 2014;14(1):14–23.

28. Lee H, Sun E, Ham D, Weissleder R. Chip–NMR biosensor for detection and molecular analysis of cells. Nature Medicine. 2008;14(8):869–874.

29. Haun JB, Devaraj NK, Hilderbrand SA, Lee H, Weissleder R. Bioorthogonal chemistry amplifies nanoparticle binding and enhances the sensitivity of cell detection. Nature Nanotechnology. 2010;5(9):660–665.

30. Issadore D, Min C, Liong M, et al. Miniature magnetic resonance system for point-of-care diagnostics. Lab on a Chip. 2011;11(13):2282.

31. Liong M, Hoang AN, Chung J, et al. Magnetic barcode assay for genetic detection of pathogens. Nature Communications. 2013;4(1):.

32. Kong TF, Peng WK, Luong TD, Nguyen N-T, Han J. Adhesive-based liquid metal radio-frequency microcoil for magnetic resonance relaxometry measurement. Lab Chip. 2012;12(2):287–294.

33. Fook Kong T, Ye W, Peng WK, et al. Enhancing malaria diagnosis through microfluidic cell enrichment and magnetic resonance relaxometry detection. Scientific Reports. 2015;5(1).

34. Kumar S, Bandyopadhyay U. Free heme toxicity and its detoxification systems in human. Toxicology Letters. 2005;157(3):175–188.

35. Çimen MYB. Free radical metabolism in human erythrocytes. Clinica Chimica Acta. 2008;390(1–2):1–11.

36. Hahn EL. Spin Echoes. Physical Review. 1950;80(4):580–594.

37. Fenimore PW, Frauenfelder H, McMahon BH, Young RD. Bulk-solvent and hydration-shell fluctuations, similar to - and -fluctuations in glasses, control protein motions and functions. Proceedings of the National Academy of Sciences. 2004;101(40):14408–14413.

38. Robinson MD, Mishra I, Deodhar S, et al. Water T2 as an early, global and practical biomarker for metabolic syndrome: an observational cross-sectional study. Journal of Translational Medicine. 2017;15(1).

39. Thulborn KR, Waterton JC, Matthews PM, Radda GK. Oxygenation dependence of the transverse relaxation time of water protons in whole blood at high field. Biochim. Biophys. Acta. 1982;714(2):265–270.

40. Gomori JM, Grossman RI, Yu-Ip C, Asakura T. NMR relaxation times of blood: dependence on field strength, oxidation state, and cell integrity. J Comput Assist Tomogr. 1987;11(4):684–690.

41. Ogawa S, Lee TM, Kay AR, Tank DW. Brain magnetic resonance imaging with contrast dependent on blood oxygenation. Proceedings of the National Academy of Sciences. 1990;87(24):9868–9872.

42. Tarburton J. Amyl Nitrite Induced Hemoglobin Oxidation Studies in Diabetics and Nondiabetics Blood. Journal of Diabetes & Metabolism. 2013;04(04).

43. Yang H, Jin X, Kei Lam CW, Yan S-K. Oxidative stress and diabetes mellitus. Clinical Chemistry and Laboratory Medicine. 2011;49(11).

44. Bourdon E, Loreau N, Blache D. Glucose and free radicals impair the antioxidant properties of serum albumin. FASEB J. 1999;13(2):233–244.

45. Roche M, Rondeau P, Singh NR, Tarnus E, Bourdon E. The antioxidant properties of serum albumin. FEBS Letters. 2008;582(13):1783–1787.

46. Lodovici M, Giovannelli L, Pitozzi V, et al. Oxidative DNA damage and plasma antioxidant capacity in type 2 diabetic patients with good and poor glycaemic control. Mutation Research/Fundamental and Molecular Mechanisms of Mutagenesis. 2008;638(1–2):98–102.

47. Grösch L, Noack F. NMR relaxation investigation of water mobility in aqueous bovine serum albumin solutions. Biochim. Biophys. Acta. 1976;453(1):218–232.

48. Cistola DP, Robinson MD. Compact NMR relaxometry of human blood and blood components. TrAC Trends in Analytical Chemistry. 2016;83:53–64.

49. Kadota K, Yui Y, Hattori R, Murohara Y, Kawai C. Decreased sulfhydryl groups of serum albumin in coronary artery disease. Jpn. Circ. J. 1991;55(10):937–941.

50. Oettl K, Stauber RE. Physiological and pathological changes in the redox state of human serum albumin critically influence its binding properties. Br. J. Pharmacol. 2007;151(5):580–590.

51. Sogami M, Era S, Nagaoka S, et al. HPLC-studies on nonmercapt-mercapt conversion of human serum albumin. Int. J. Pept. Protein Res. 1985;25(4):398–402.

52. Sun N, Yoon T-J, Lee H, et al. Palm NMR and 1-Chip NMR. IEEE Journal of Solid-State Circuits. 2011;46(1):342–352.

53. Haun JB, Castro CM, Wang R, et al. Micro-NMR for Rapid Molecular Analysis of Human Tumor Samples. Science Translational Medicine. 2011;3(71):71ra16–71ra16.

54. Genuth S, Sun W, Cleary P, et al. Glycation and carboxymethyllysine levels in skin collagen predict the risk of future 10-year progression of diabetic retinopathy and nephropathy in the diabetes control and complications trial and epidemiology of diabetes interventions and complications participants with type 1 diabetes. Diabetes. 2005;54(11):3103–3111.

55. Lee R, Margaritis M, Channon KM, Antoniades C. Evaluating oxidative stress in human cardiovascular disease: methodological aspects and considerations. Curr. Med. Chem. 2012;19(16):2504–2520.

